# Cytosol-living *Trypanosoma cruzi* amastigotes scavenge cholesterol from host ER and Golgi complex

**DOI:** 10.1101/2024.07.10.602841

**Authors:** Carolina de Lima Alcantara, Miria Gomes Pereira, Wanderley de Souza, Narcisa Leal da Cunha-e-Silva

**Affiliations:** Laboratório de Ultraestrutura Celular Hertha Meyer, Instituto de Biofísica Carlos Chagas Filho, Universidade Federal do Rio de Janeiro, Rio de Janeiro, Brazil; Centro Nacional de Biologia Estrutural e Bioimagem, Universidade Federal do Rio de Janeiro, Rio de Janeiro, Brazil

## Abstract

Chagas Disease, caused by the protozoan parasite *Trypanosoma cruzi,* stands as a prevalent neglected disease in Latin America. The amastigote stage, the replicant intracellular form of the parasite, is a crucial player in infection persistence within vertebrate hosts. The amastigotes exhibit remarkable adaptability regarding the cell type that they infect, being able to modulate their metabolism and growth based on the host cell resources availability. Lipid metabolism emerges as a key determinant of amastigote growth, with a dependency on the host cell’s lipid resources. While the parasite can synthesize some sterols and fatty acids, it also scavenges these essential components, particularly cholesterol, from the host. Alterations in the host’s cholesterol metabolism, potentially regulated by SREBPs, contribute to increased intracellular cholesterol levels, fostering parasite development. However, the mechanisms underlying cholesterol uptake by amastigotes remained elusive. Here, we investigate the cholesterol trafficking mechanism from host cells to amastigotes by employing a fluorescent cholesterol analog. Using advanced imaging techniques, such as confocal fluorescence microscopy and high-resolution volume electron microscopy we demonstrated that amastigotes internalize extracellular-derived cholesterol, defined the uptake kinetics for cholesterol by amastigotes, and demonstrated that cholesterol is important for amastigote development. Analysis of the interaction of host cell ER with the amastigotes revealed by the presence of membrane contact sites between this organelle and the amastigote plasma membrane. We also showed that amastigotes can take up host ER and Golgi proteins, probably by endocytosis, paving a new mechanism for host cell scavenging of molecules by the parasite.

**Author Summary:** Chagas Disease, a widespread yet often overlooked challenge in Latin America, is driven by the parasite *Trypanosoma cruzi*. Understanding the amastigote stage, the parasite’s replicative form, is crucial for grasping infection persistence. This study reveals a critical survival strategy – amastigotes adeptly scavenging cholesterol from host cells. We demonstrated that the amastigotes can take up cholesterol from the host and that cholesterol is important for parasite development. Analysis of host cell organelles involved with cholesterol traffic, ER and Golgi, suggested their involvement, establishing crucial conduits for cholesterol acquisition. Deciphering these mechanisms illuminates the intricate interplay between parasite and host, presenting a potential treatment target. The parasite’s reliance on host cholesterol underscores adaptable growth strategies. This breakthrough deepens Chagas Disease understanding, paving the way for targeted therapeutic interventions.

## Introduction

Chagas Disease is one of the most prevalent neglected diseases in Latin America [1] and is caused by infection with the protozoan parasite *Trypanosoma cruzi*. *T. cruzi* life cycle takes place in vertebrate and invertebrate hosts and comprises proliferative and infective forms that are highly adaptable to their hosts to maintain the infection. In the vertebrate host, amastigote forms are responsible for parasite replication and persistence of the infection in chronically infected patients. Amastigotes live free in the host cell cytosol having access to a myriad of macromolecules from the host and host cell organelles. Scavenge of nutrients from hosts is central to a parasitic lifestyle. One of the parasites’ major mechanisms of nutrient salvage is the presence of plasma membrane transporter proteins [2]. Some works have suggested the participation of transporters at the cell surface of amastigotes that may help in micronutrient and ion acquisition to sustain their growth and replication [3–5]. However, amastigotes also possess an active machinery for endocytosis: a specialized membrane invagination called cytostome-cytopharynx complex through which macromolecules are endocytosed [6,7]. It has been already shown that amastigotes recently released from the host cell cytosol by mechanical disruption can endocytose albumin and transferrin added to the extracellular incubation medium [6,8]. Labeling cytosolic host proteins with a fluorescent probe showed that amastigotes cannot do bulk endocytosis of cytosolic host proteins, suggesting that other mechanisms control the endocytosis of host cell components [6].

Although many groups are involved in understanding how the parasite manipulates the cell traffic for its benefit, the lipid traffic between the host towards amastigotes is slightly comprehended. Besides, some amastigotes are found in dormancy during the chronic stage [9,10], therefore, more pharmacological strategies are needed to control parasite proliferation. The lipid routes arose as an alternative to Chagas disease treatment.

Biochemical studies have shown that *T. cruzi* epimastigotes, the replicative forms found in the insect mid-gut, can synthesize de novo its own sterols, mainly ergostane-type sterols, as well as scavenging cholesterol from the host [11] via endocytosis of lipoproteins. Epimastigotes grown in axenic culture can endocytose LDL and accumulate the released cholesterol in lipid inclusions in the reservosomes or lipid droplets in the cytosol, being able to mobilize these stocks when necessary [12–14].

Amastigotes are also able to synthesize *de novo* sterols (ergosta-7-en-3β-ol, ergosta-7,24(24^1^)- dien-3β-ol (episterol) or 24-ethyl-7-en-cholesta-3β-ol) [15], and fatty acids [16], and is also capable to scavenge sterols from the mammalian host, evidenced by the presence of cholesterol and cholesta-5,24-dien-3β-ol (desmosterol) [15]. These studies have shown that cholesterol corresponds to 80% by weight of total sterols in amastigotes.

Omics works in the last decade have provided important clues about host cell and parasite cellular pathways modulated during infection [16–18]. In this context, it has been shown that amastigote growth and replication are highly dependent on the lipid metabolism of the host cell [17,18]. *T. cruzi* infection can lead to changes in the cholesterol metabolism of the host, by increasing intracellular cholesterol levels [19], probably by upregulation of the SREBPs (sterol regulatory element binding proteins) that regulate lipid homeostasis [16], favoring parasite development. However, direct evidence for the role of cholesterol for parasite growth and development is absent.

Despite the biochemical evidence pointing to a role of host lipids in the metabolism of intracellular amastigotes, the mechanisms by which amastigote scavenge these lipids are unclear. In this work, using a fluorescent cholesterol analog, we could follow cholesterol traffic from the host cells to the amastigotes using confocal fluorescence microscopy. We were able to define the kinetics of cholesterol acquisition by amastigotes and the possible host organelles involved in the traffic. Moreover, we demonstrated that cholesterol is important for amastigote grown and development. Ultrastructural analysis by electron tomography and FIB-SEM (*<u>F</u>ocused Ion <u>B</u>eam-<u>S</u>canning <u>E</u>lectron <u>M</u>icroscopy*) showed he the presence of contact sites between the parasite and host ER and Golgi complex may be responsible for the acquisition of cholesterol by amastigotes and provided important information about these organelles remodeling in the course of *T. cruzi* infection.

## Results

### Amastigotes internalize extracellular-derived cholesterol

To follow the traffic of cholesterol from the host cell to the amastigotes, we first used LDL particles loaded with a fluorescent cholesterol analog, TopFluor Cholesterol (TopFChol) diluted in delipidated FBS (dFBS), to favor cholesterol acquisition by the cells. Infected host cells were incubated with LDL-TopFChol for different times: a short incubation of 4h and a longer incubation of 24h. As expected, after 4h of incubation, LDL-TopFChol signal was observed mainly in LAMP1 positive structures (Fig 1a), showing that LDL particles were internalized by classical receptor-mediated endocytosis and ended up in the host lysosomes. At this time point no major colocalization of the TopFChol signal with the host ER was observed (Fig 1b). At this short incubation time, it was already possible to see punctual labeling of the TopFChol inside amastigotes, especially at the anterior region of the parasites (Figs 1a and 1b). After 24h of incubation, the TopFChol signal derived from the endocytosed LDL no longer had a strong colocalization with the lysosomes (Fig 1c). Nevertheless, it showed a spread to the host cell ER (Fig 1d), agreeing with has been characterized for LDL-derived cholesterol traffic in mammalian cells (reviewed in [20]). At this point, more amastigotes showed spotted labeling inside their cytosol, mostly at the parasités anterior region, close to the main endocytic portals, the flagellar pocket (FP), and the cytostome-cytopharynx complex (Fig 1c-d). Of note, we observed labeling for the luminal ER protein PDI (*<u>P</u>rotein <u>D</u>isulfide Isomerase*) inside amastigotes (Fig 1b).

**Fig 1.**
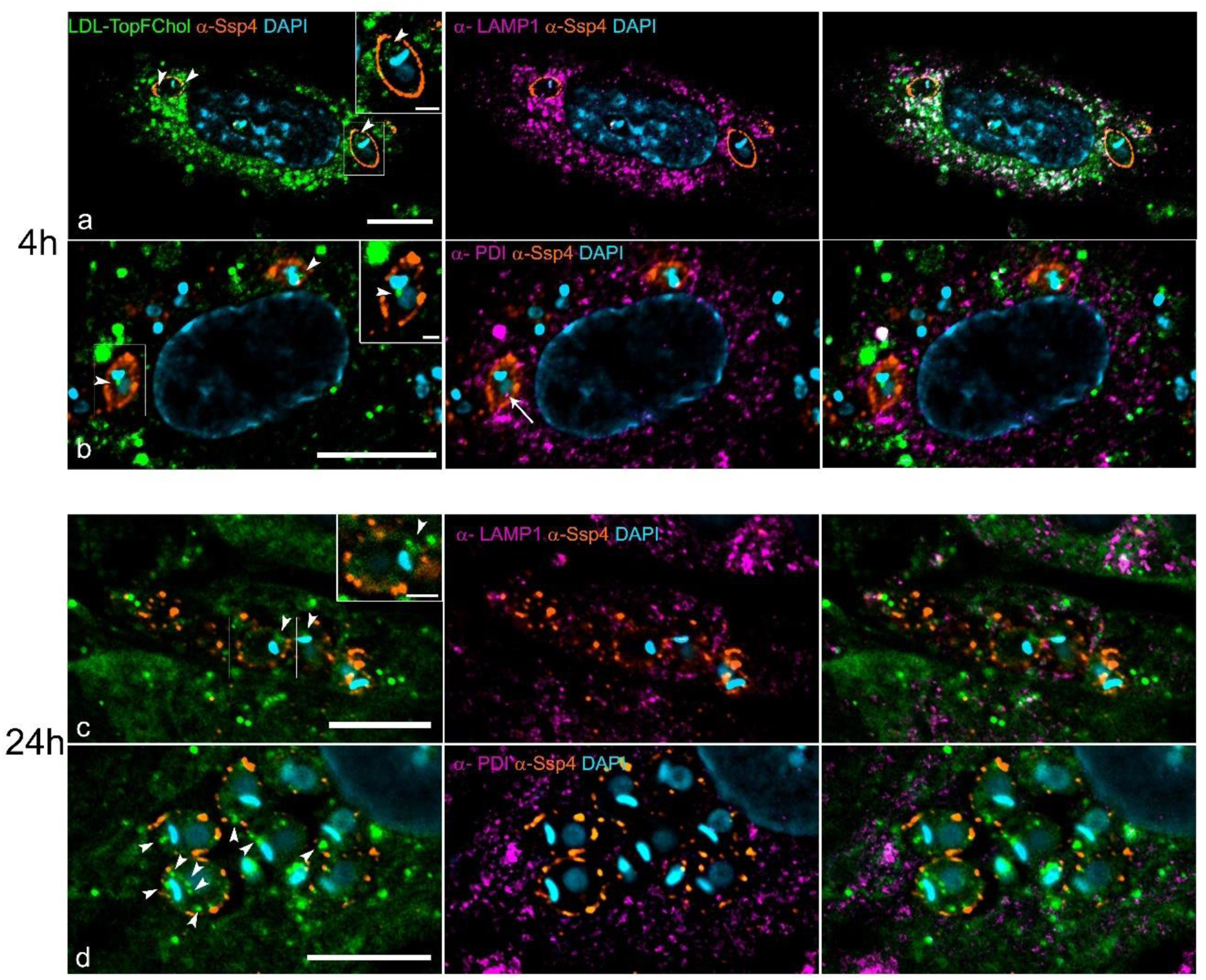
Dynamics of TopFChol internalization by intracellular amastigotes through incubation with LDL-TopFChol. HFF1 cells were infected with trypomastigotes and, 24h after, incubated with LDL-TopFChol in a growth medium supplemented with 10% of dFBS. Single-plane fluorescence images from the z-stack confocal series are shown. (a,b) Cells were incubated with LDL-TopFChol (green) for 4h, fixed and co-stained with anti-Lamp1 (to label lysosomes) or anti-PDI (to label ER) antibodies (magenta). TopFChol signal colocalized with anti-Lamp1 staining (a) but not with anti-PDI (b). Intracellular TopFChol labeling was observed inside amastigotes in punctual locations at the cell anterior (a) and posterior to the kinetoplast (b) (arrowheads in insets). Anti-Ssp4 antibody was used to label the amastigote cell membrane (orange) and DAPI (blue) stained cells DNAs. Of note, an anti-PDI signal was observed inside amastigotes at 4h at the post-nuclear location (b, arrow). (c,d) LDL-TopFChol was incubated with infected cells for 24h. TopFChol signal no longer colocalizes with anti-Lamp1 labeling (c), instead, showed a spread labeling through the cytoplasm, colocalizing with the anti-PDI signal at the ER (d). At this time point, more amastigotes showed intense punctual intracellular labeling for TopFChol (arrowheads). Bars: 10 µm.

We investigated other routes of cholesterol traffic into the host cell by incubating the TopFChol directly into the culture medium. In this condition, most cholesterol molecules should be inserted into the plasma membrane (PM) where they can be internalized through endocytosis or sequestered into the ER via PM-ER contact sites, restoring the cholesterol levels at the PM [20]. We observed that, after 4h of incubation, several amastigotes already showed spots of TopFChol inside the parasites, preferably at the anterior region (Fig 2a and 2b). After 24h of incubation, a higher intensity labeling of the TopFChol was observed inside the amastigotes (Fig 2c). Here, we also observed labeling for PDI ER protein inside amastigotes in locations that could or could not colocalize with TopFChol signal.

**Fig 2.**
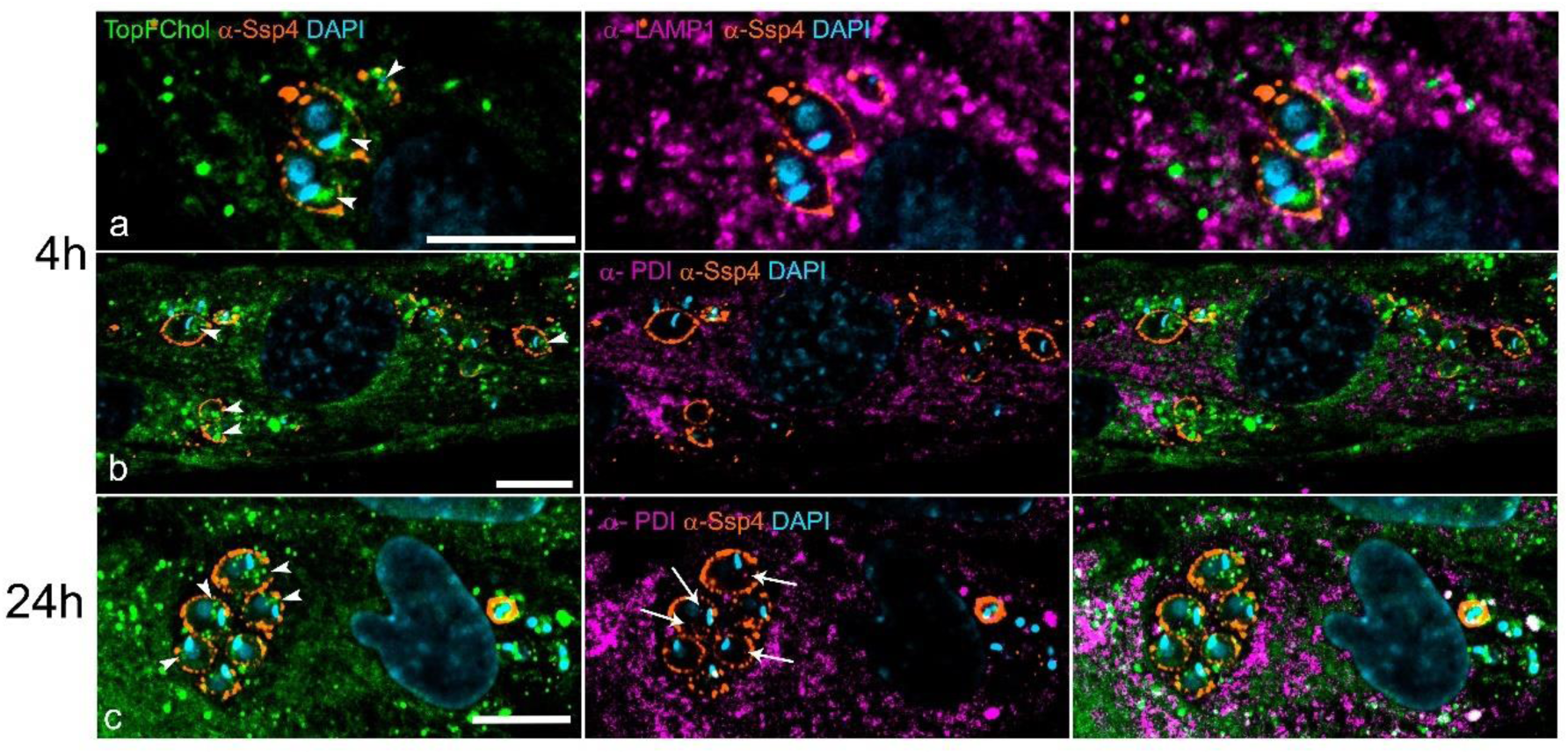
Dynamics of TopFChol internalization by intracellular amastigotes through incubation with TopFChol directly into the medium. HFF1 cells were infected with trypomastigotes and, 24h after, TopFChol was added directly to the medium supplemented with 10% of dFBS. Single-plane fluorescence images from the z-stack confocal series are shown. (a,b) Cells were incubated with TopFChol (green) for 4h, fixed and co-stained with anti-Lamp1 (a) or anti-PDI (b) antibodies (magenta). An intense TopFChol signal can be observed inside the amastigotes (arrowheads in a,b). No colocalization of TopFChol signal and Lamp1-positive compartments was observed (a), but, instead seemed to colocalize with PDI-positive compartments (b). (c) TopFChol was incubated with infected cells for 24h. Amastigotes showed intense punctual TopFChol labeling along the anterior-posterior region (arrowheads). Labeling for PDI was observed inside amastigotes in regions that colocalized with the TopFChol signal (arrow). Anti-Ssp4 antibody was used to label the amastigote cell membrane (orange) and DAPI (blue) stained cells DNAs. Bars: 10 µm.

The pattern of labeling of the TopFChol inside the amastigotes was peculiar, usually forming a winding or helical shape from the cell’s anterior region, where it was more concentrated, to the cell’s posterior (S1 Fig a). Using immunogold labeling with silver enhancement for the BODIPY moiety of TopFChol in LRWhite sections, we observed an intense labeling at the preoral ridge (POR), flagellar pocket and in spots along the amastigote plasma membrane, in infected cells incubated with TopFChol for 24h (S1 Figs b-d). Silver enhanced labeling was stronger in areas of the host cell, besides the parasite membrane. We also observed labeling at the membrane of the cytopharynx of an amastigote (S1 Fig d). Next, we quantified the presence or absence of TopFChol labeling in amastigotes and the location of the signal inside them (anterior or posterior) at the different times and availability of TopFChol. For that, we divided the amastigotes into two halves, i.e anterior, from the flagellum tip until the posterior portion of the kinetoplast, and posterior, from the perinuclear region down to the posterior tip (Fig 3a). Two hundred amastigotes were counted for each condition. After 4h, in cells incubated with TopFChol added directly to the culture medium, a mean of 72% of the amastigotes presented intracellular punctual labeling, both at celĺs anterior and posterior, when compared to only 4% of the amastigotes in host cells incubated with TopFChol loaded into LDL particles (Fig 3b). After 24h, a mean of 77% of the amastigotes presented labeling of TopFChol both at cell’s anterior and posterior when host cells were incubated with TopFChol in the medium while a mean of only 18% present this labeling pattern when host cells were incubated with LDL-TopFChol. This indicated that Chol was internalized by the amastigotes, in both conditions. However, the traffic route of the tracer inserted into the PM, by adding TopFChol directly to the culture medium, was faster than when Chol was provided inside the LDL particles. Next, we evaluated if cholesterol was important for amastigote proliferation. For that, we incubated the infected cells under three different conditions: medium supplemented with 10% whole FBS (control), or with 10% dFBS or with 10%FBS + TopFChol (Fig 3c). Infected cells and the number of amastigotes/cells were evaluated 48hpi. We observed a decrease in both infectivity and number of amastigotes/cell when cells were incubated in dFBS, compared with control. However, infectivity and amastigotes/cell were restored to control if TopFChol was added to the medium with dFBS (Fig 3d). This result demonstrated that amastigote proliferation was sensitive to a decrease in cholesterol availability.

**Fig 3.**
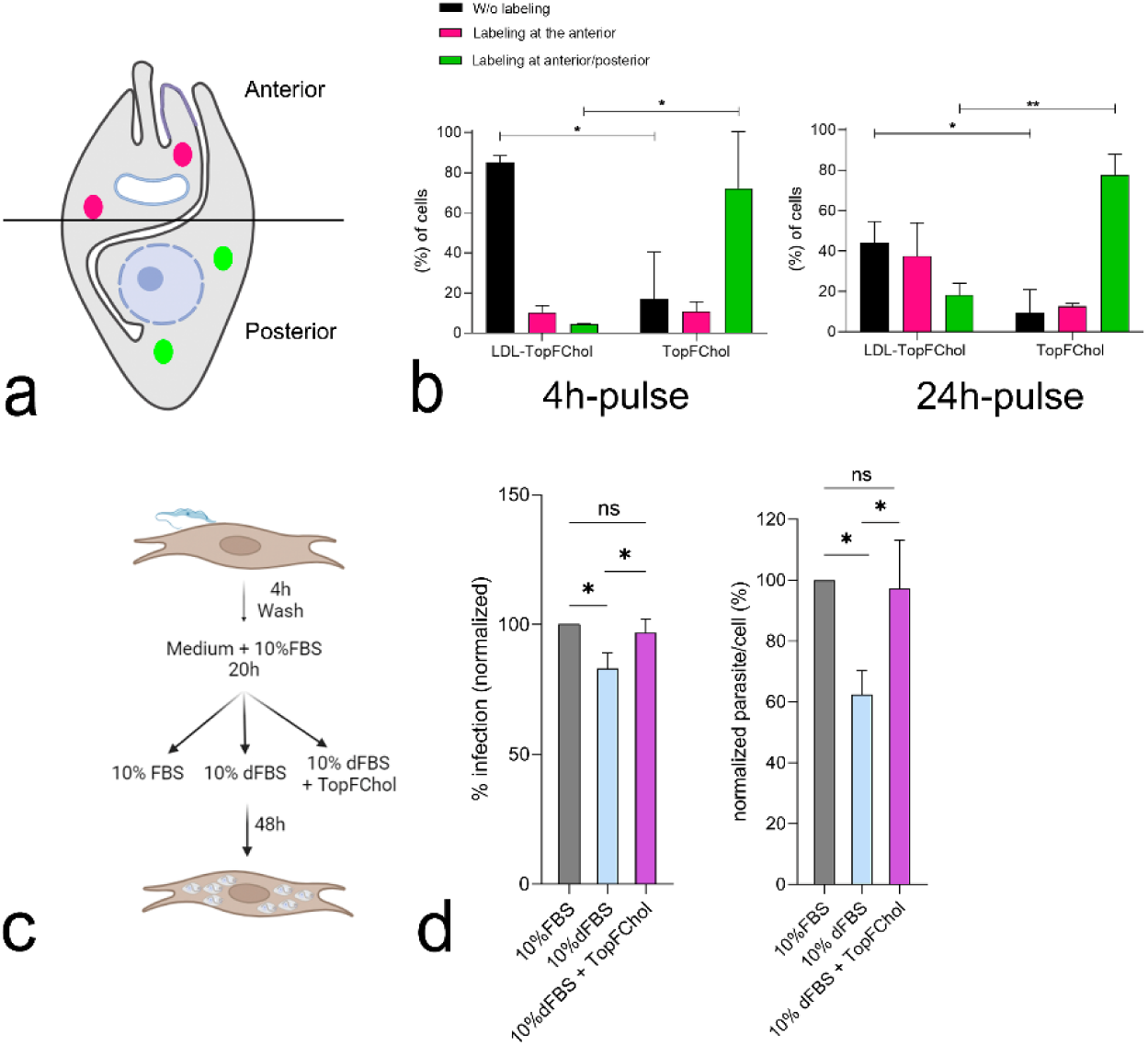
Kinetics of TopFChol internalization by amastigotes and cholesterol effect in amastigote proliferation. (a) Schematic representation of the amastigote division parameter used for the analysis shown in b. The amastigote cell body was divided into two halves. All labeling localized from the anterior region until the posterior of the kinetoplast was considered to be at the cell’s anterior halves. All labeling localized from the posterior of the kinetoplast to the posterior tip of the amastigotes was tagged as posterior labeling. (b) HFF1-infected cells were incubated with LDL-TopFChol or TopFChol directly in the culture medium for 4 or 24h. Confocal z-stacks were acquired and TopFChol labeling inside amastigotes was quantified using the criterion shown in d. 200 amastigotes were counted in each condition. Data represent mean ± SD values. *p < 0.01 and **p < 0.001 analyzed by two-way ANOVA followed by Bonferroni. (c) Schematic representation of the experimental design used for determination of amastigotes proliferation in different availability of lipids. (d) Infection efficiency and the numbers of amastigote per infected cell were determined by counting a total of 300 HFF cells (n=2 in triplicate) in each condition. Data was normalized by the control (10%FBS). Mean±sd are shown. Data were analyzed using One-way ANOVA with Post Hoc Holm-Šídák’s multiple comparisons test

### Role of host biosynthetic-secretory organelles in cholesterol scavenge by amastigotes

#### Amastigote-ER contact sites

The results shown so far point to a probable role of host ER in the traffic of cholesterol for the amastigotes. As mentioned, we observed antibody labeling for host PDI, an ER luminal protein, in punctual locations inside amastigotes. To rule out the possibility of cross-reactivity of the antibody with amastigote’s antigens, we used a transient transfection system, Cell Light ER-RFP, that uses a baculovirus vector to deliver a fusion construct of ER signal sequence of calreticulin and KDEL (ER retention signal) and the fluorescent tag RFP. Infected cells were incubated overnight with the reagent and the next day incubated with LDL-TopFChol (S2 Fig) or TopFChol directly in the culture medium (S3 Fig). In both cases, we observed the same labeling kinetics as before, with 4h and 24h of incubation, with a faster delivery of TopFChol to amastigote when TopFChol was added to the culture medium. However, we observed stronger labeling of the Cell Light ER-RFP inside the amastigotes that colocalized with the TopFChol signal, especially at 24h (S3 Fig). These results provided strong evidence that, at least some, host-ER proteins can be internalized by the amastigotes. We extended the incubation time to 48h and saw a change in the Cell Light ER-RFP labeling from reticulated to a more punctate staining pattern, inside and around the amastigotes, whatever TopFChol was provided inside LDL or free in the culture medium (S2 and S3 Figs).

We then analyzed, by TEM, the ultrastructure of infected cells incubated under normal culture conditions, i.e. medium with 10% FBS (Figs 4a-d) or under lipid deprivation, i.e. medium with 10% delipidated FBS (Figs. 4e-i). In the whole serum, lipids are available for endocytosis by the host cells. In delipidated serum, host cells may activate endogenous cholesterol biosynthesis at ER. In cells, 48hpi, incubated with whole serum, close contact of host lysosomes and ER with the amastigote’s membrane was observed. The juxtaposition of these organelle’s membranes and amastigote membrane was so close that it was difficult to distinguish their boundaries (Figs 4a-c). Of note, some cytosolic material and ER profiles could be seen at the entrance of the amastigotés cytostome and in contact with the POR membrane domain (Figs 4c and 4d). On the other hand, in cells incubated with dFBS, ER was the most prominent organelle surrounding the amastigotes (Fig 4e). Many contact sites could be observed between ER and the amastigote membrane, especially at the anterior region of the parasite (Figs 4f-i).

**Fig 4.**
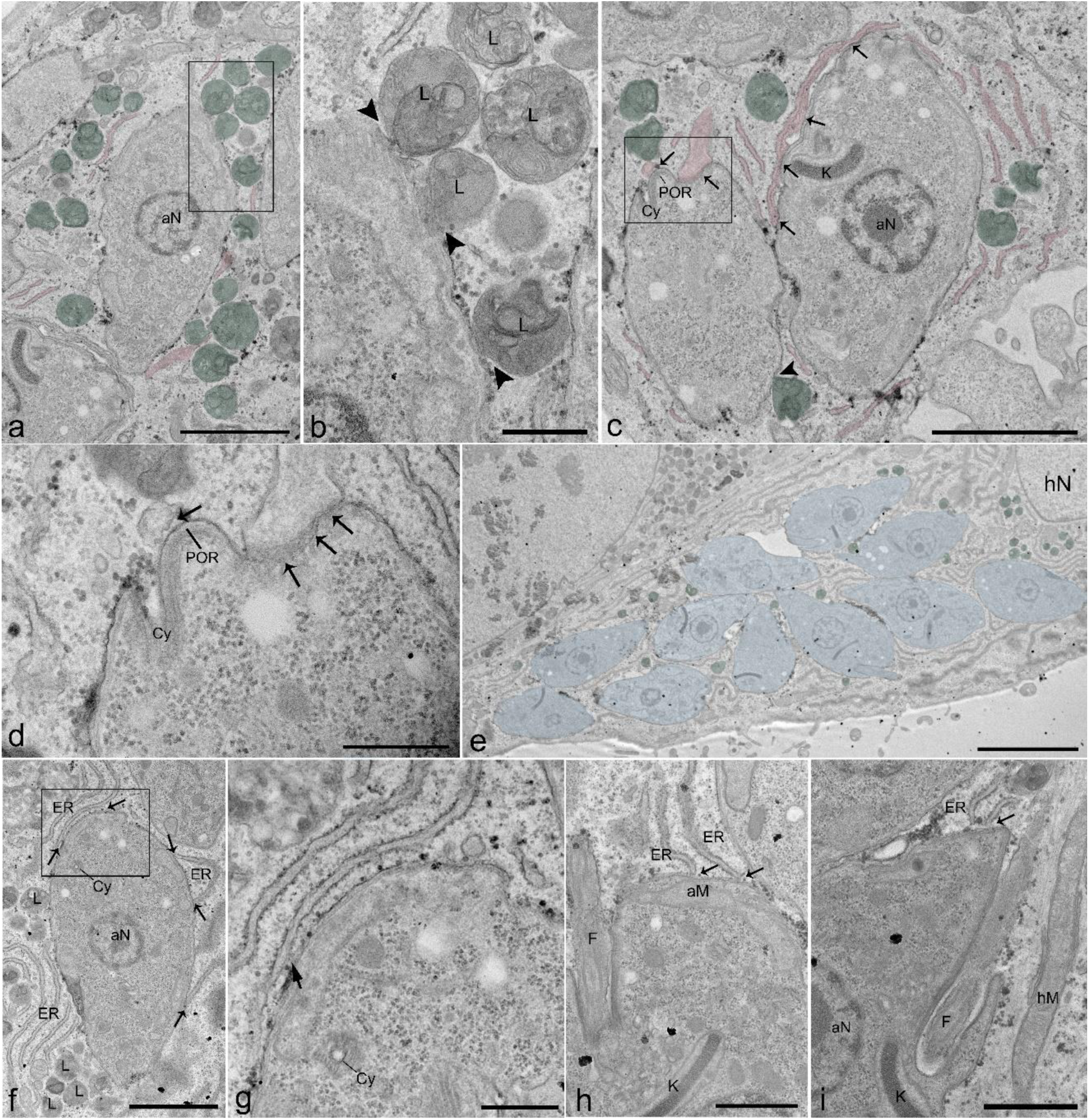
Differences in host cell distribution around amastigotes in different conditions of lipids availability. HFF1 cells were infected with trypomastigotes and, 24h after, the growth medium was changed, and cells were incubated with 10% of complete FBS (control) or with 10% dFBS (lipid deprivation). Cells were fixed 48h later and processed for TEM. (a-d) TEM images of cells incubated with complete FBS. (a) Host lysosomes (L, green) and ER (pink) could be seen very close to the amastigote membrane. (b) Higher magnification of the region highlighted by the rectangle in a. Lysosomes showed direct contact with the amastigote membrane (arrowhead). (c) ER appeared to embrace amastigotes at some points of contact (arrows). (d) Higher magnification of the region highlighted by the rectangle in c. ER profiles could be seen touching the amastigote membrane at the region of the preoral ridge (POR) and the cytostome (Cy) (arrows). (e-i) TEM images of cells incubated with 10% dFBS. (e) Low magnification TEM image showing an infected cell full of amastigotes (blue), prominent ER, and a few lysosomes (green). (f-i) ER profiles could be seen touching the membrane of amastigotes (arrows) at many points, especially at the amastigote anterior region (g-i). aN (amastigote nucleus); hN (host nucleus); aM (amastigote mitochondria); hM (host mitochondria); F (flagellum); K (kinetoplast). Bars: a,c,f - 2 µm; b,d,g – 500 nm; e - 5 µm; h, i - 1 µm.

Transmission electron microscopy has long been acknowledged as the benchmark method for observing the intricate details of cellular ultrastructure. However, bi-dimensional images lack the volume and cellular context needed to describe intraorganellar membrane contact sites (MCS), which requires higher-resolution microscopy methods such as electron tomography. We then analyzed tomograms acquired from 200nm sections from 48hpi cells cultivated with 10% of whole serum. In total, 11 tomograms (pixel size of 1.94 nm) were acquired and analyzed. As seen previously in Fig.4, we observed an intimate contact of host ER and lysosome membranes with the amastigote plasma membrane (Figs 5a and 5b). Bridges connecting host lysosomes and amastigotes (Fig 5c) and host ER and amastigotes (Fig 5d) were clearly identified in the tomograms. Concerning ER-amastigotes MCSs, bridges of different extensions could be observed. Some were very short, bringing ER and amastigote to juxtaposition (Fig 5d), and some were longer and seemed to tether both membranes (Fig 5e-i). We measured the number of bridges between the amastigote membrane and the main organelles that appeared around the parasites in the tomograms, i.e. ER, lysosomes, and mitochondria. Plotting this measurement (Fig 5j) we could demonstrate that ER was the organelle that formed more bridges with the amastigote membrane, with an average of 10 bridges per tomogram. We also measured the length of the bridges between the amastigote membrane and the host organelles (Fig 5k). Most bridges between hER-amastigote varied from 5-30 nm but longer bridges could reach 80 nm. hMito-amastigote bridges were more homogeneous, varying between 5-40 nm, and hLysosome-amastigote bridges were restricted to lengths varied from 6-10 nm.

**Fig 5.**
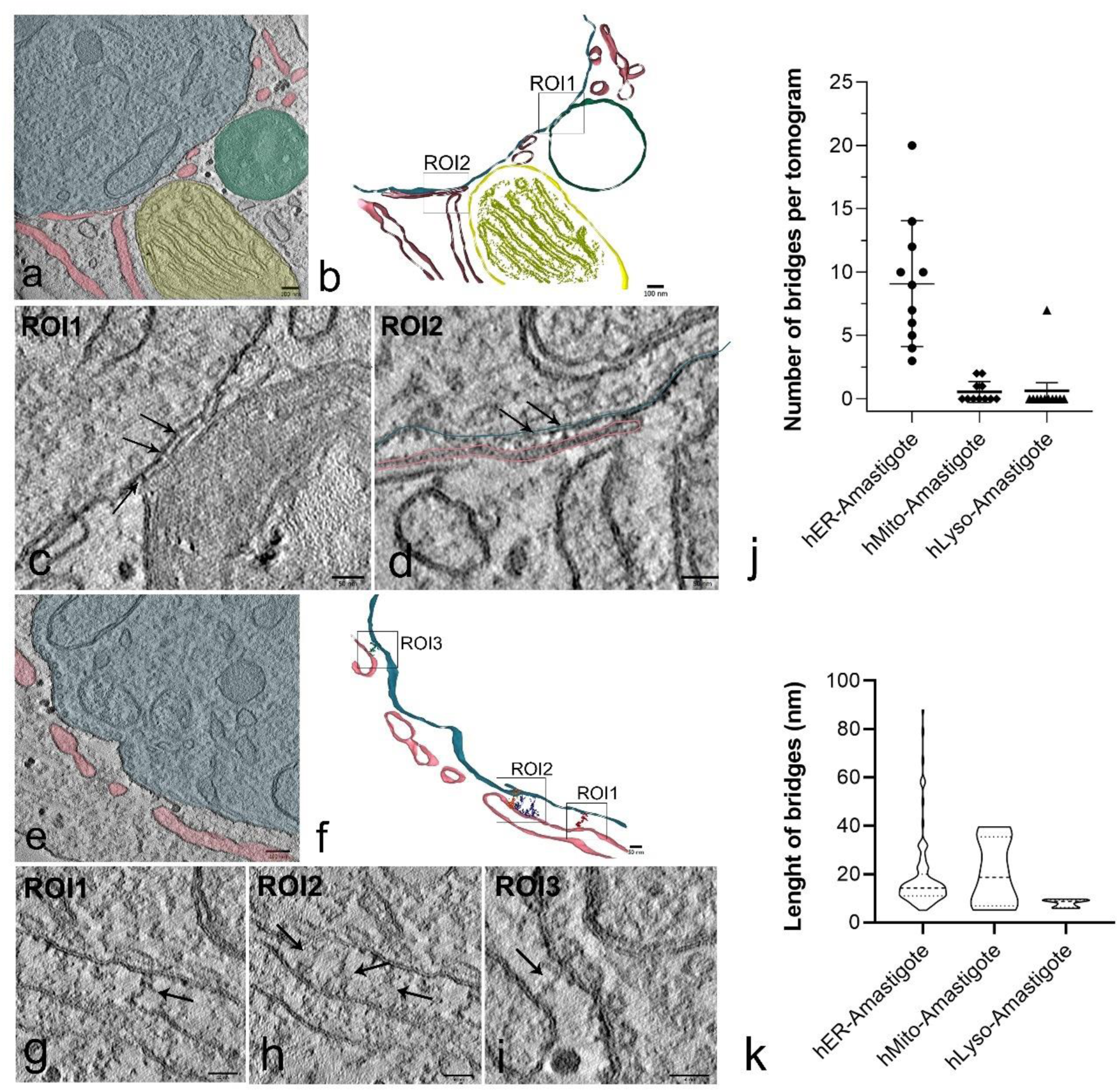
Host organelles exhibit contact sites with amastigote membrane. HFF1 cells were infected with trypomastigotes and incubated with the growth medium supplemented with 10% of complete FBS (control) for 48h. Cells were then fixed and processed for TEM. (a) A tomogram virtual slice image showing the main host organelles found in the vicinity of amastigotes (colored in blue): host ER (pink), mitochondria (yellow), and lysosomes (green). Note the proximity of the ER membrane and lysosome membrane to the amastigote. (b) 3D reconstruction of the tomogram showed in a, highlighting regions of intimate contact between lysosome-amastigotes (ROI1) and ER-amastigotes (ROI2). Tomogram images of these ROIs are shown in c and d, respectively. Electron-dense bridges connecting host lysosome membrane (c) and host ER membrane (d) with the amastigote membrane could be observed (arrows). (e) Virtual slice image of another tomogram showing ER profiles (pink) in the vicinity of the amastigote membrane (blue). In the 3D reconstruction shown in f, bridges between the two membranes were segmented by threshold and are highlighted in ROIs 1,2 and 3. These same regions can be observed in the tomogram images shown in g,h, and f, where the bridges are pointed by the arrows. (j) Measure of the number of bridges between each host organelle and amastigote membrane per tomogram (n=11). (k) Measure of the length of the bridges between host organelles and amastigotes. The graph shows the frequency distribution of the bridge’s lengths found for each organelle. Bars: a, b, e – 100nm; c,d,f-1 – 50 nm.

#### Role of host Golgi complex in cholesterol acquisition by the amastigotes

Cholesterol homeostasis in the cells is strictly controlled by the ER, and the cholesterol gradient ranges from low concentration at the ER to higher concentration at the PM. Cholesterol at the ER is transported to the Golgi via non-vesicular transport and is enriched at the trans-Golgi [20], from where any vesicles bud off as part of the secretory pathway. Based on the role of Golgi apparatus in the secretion of molecules, we also decided to analyzed its contribution in cholesterol traffic to the amastigotes.

Based on our previous results, incubation of cells with TopFChol directly into the culture medium was the faster way for cholesterol to reach the amastigotes. In this context, cholesterol should be sequestered from PM by the ER and then transferred to the Golgi complex. Based on this, we incubated infected cells with TopFChol for 24h and immunolabelled the host Golgi complex with an antibody that recognized a resident integral membrane protein of the trans-Golgi network, TGN-38 [21]. We observed dispersed punctual labeling of anti-TGN38 antibody localized at the cell cytosol that colocalized with the TopFChol stain (Figs 6a and 6b). Also, labeling for TGN38 was observed inside the amastigotes colocalizing with TopFChol, especially at the amastigote anterior region. Again, to rule out the cross-reactivity of the antibody with amastigote antigens, we performed the same experiment using the reagent Cell Light Golgi-RFP. In this case, the construct encodes a human trans-Golgi resident enzyme, N-acetylgalactosaminyltransferase, tagged with RFP. We also observed labeling of the Cell Light reagent inside amastigotes, colocalizing with the TopFChol signal (Figs 6 c and 6d). These results showed that, like ER, the Golgi complex also participated in the traffic of cholesterol to the amastigotes and that the parasite somehow could acquire proteins from the Golgi as well. In infected cultures incubated with TopFChol for 24 and processed for TEM, images of thin sections showed the proximity of the Golgi cisternae with the anterior region of the amastigote cell membrane. We also observed several small Golgi stacks spread to the cell cytoplasm (Figs 6 e and 6f).

**Fig 6.**
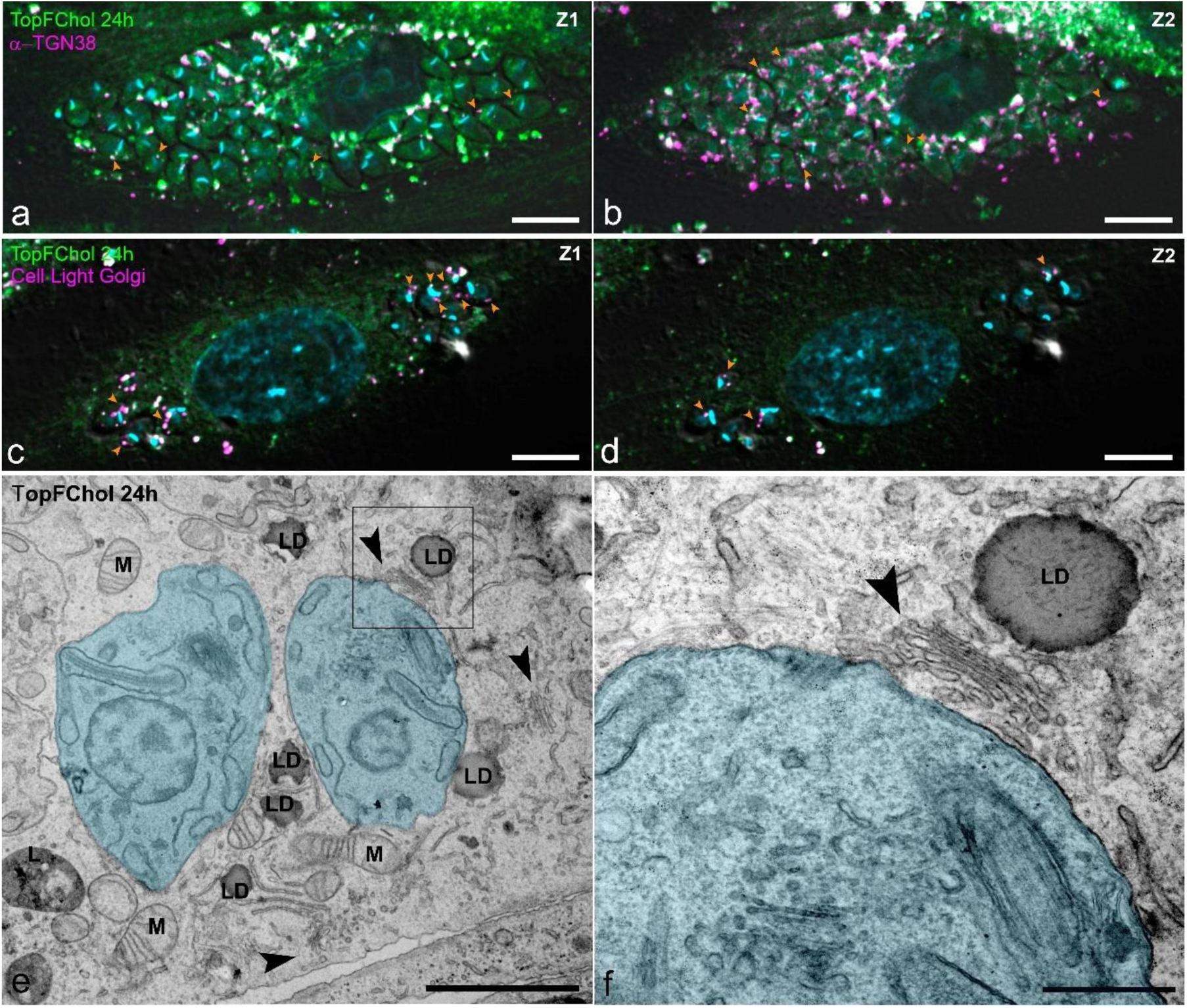
Host Golgi complex participates in TopFChol delivery to amastigotes. HFF1 cells were infected with trypomastigotes and, 24h after, TopFChol was added directly to the growth medium supplemented with 10% of dFBS, for 24h. Cells were fixed and immunolabeled with antibody anti-TGN38 (a,b), or cells were incubated for 24h with Cell Light Golgi-RFP before fixation (c,d). (a,b) Sequential planes of fluorescence images from the z-stack confocal series. (a) TopFChol signal (green) could be seen labeling the amastigote membrane (note that it was possible to delimitate the amastigote body) and also in spots at the amastigote cell interior (arrowheads). (b) in the sequential plane, labeling for the TGN-38 (magenta) could be found colocalizing with TopFChol inside the amastigotes (arrowheads). The colocalization between the TopFChol signal and Cell Ligh Golgi was also seen inside the amastigotes (c,d). In infected cells processed for TEM, we observed the presence of Golgi stacks dispersed through the cytosol (arrowheads in e), some of them lining very close to the amastigote membrane (arrowhead in f, from the rectangle region highlighted in e). M (Mitochondria), LD (Lipid droplets), L (Lysosomes). Bars: a-d – 10 µm; e - 2 µm; f – 500 nm.

In vertebrate cells, the Golgi apparatus is arranged in multiple stacks that associate laterally forming the Golgi ribbon, located perinuclearly [22,23]. As we observed scattered stacks in cells incubated with TopFChol, we analyzed the organization of the Golgi complex in host cells after infection and different availability of cholesterol. For that, the Golgi complex was labeled with anti-TGN38 antibody in non-infected cells and infected cells 24hpi cultivated with 10% FBS or 10% dFBS (S4 Fig). In non-infected cells, the antibody labels the Golgi similarly with the already described perinuclear ribbon (S4 Fig a). In infected cells cultivated with 10% FBS, the antibody labeled small spots disperse through the cell cytosol (S4 Fig b). Signal of the antibody could be seen inside the amastigotes (S4 Fig b-inset), with an accumulation at the amastigote anterior region and spots that spanned from the anterior to the posterior of the parasite. Infected cells incubated with 10% dFBS, showed more intense labeling of the Golgi, with several dispersed spots throughout the cytosol (S4 Fig c). In this case, we also observed labeling of the antibody inside the amastigotes, following a path from the parasite anterior, where it was more concentrated, to the parasite posterior region (S4 Fig c-inset). This result shows that the structural organization of the Golgi apparatus in the host cells is modified after infection, with Golgi fragmentation in small stacks and dispersion through the cytosol, facilitating its contact with the amastigotes.

By using serial electron tomography and FIB-SEM, we were able to get a tridimensional overview of the interaction between the Golgi stacks and amastigotes. We acquired 3 serial tomograms (4 sections of 200 nm each – pixel size: 1.94 nm) of infected cells 48hpi incubated in culture medium supplemented with 10% FBS. The tomogram shown in Fig 7a, reveals the proximity of the host Golgi stack to the anterior region of amastigotes, close to the flagellar pocket and the preoral ridge (POR). The 3D reconstruction of the tomogram volume shows the distribution and proximity of the host ER to the amastigote membrane, that surrounds the amastigote entirely (Figs 7b and 7c and Movie 1).

**Fig 7.**
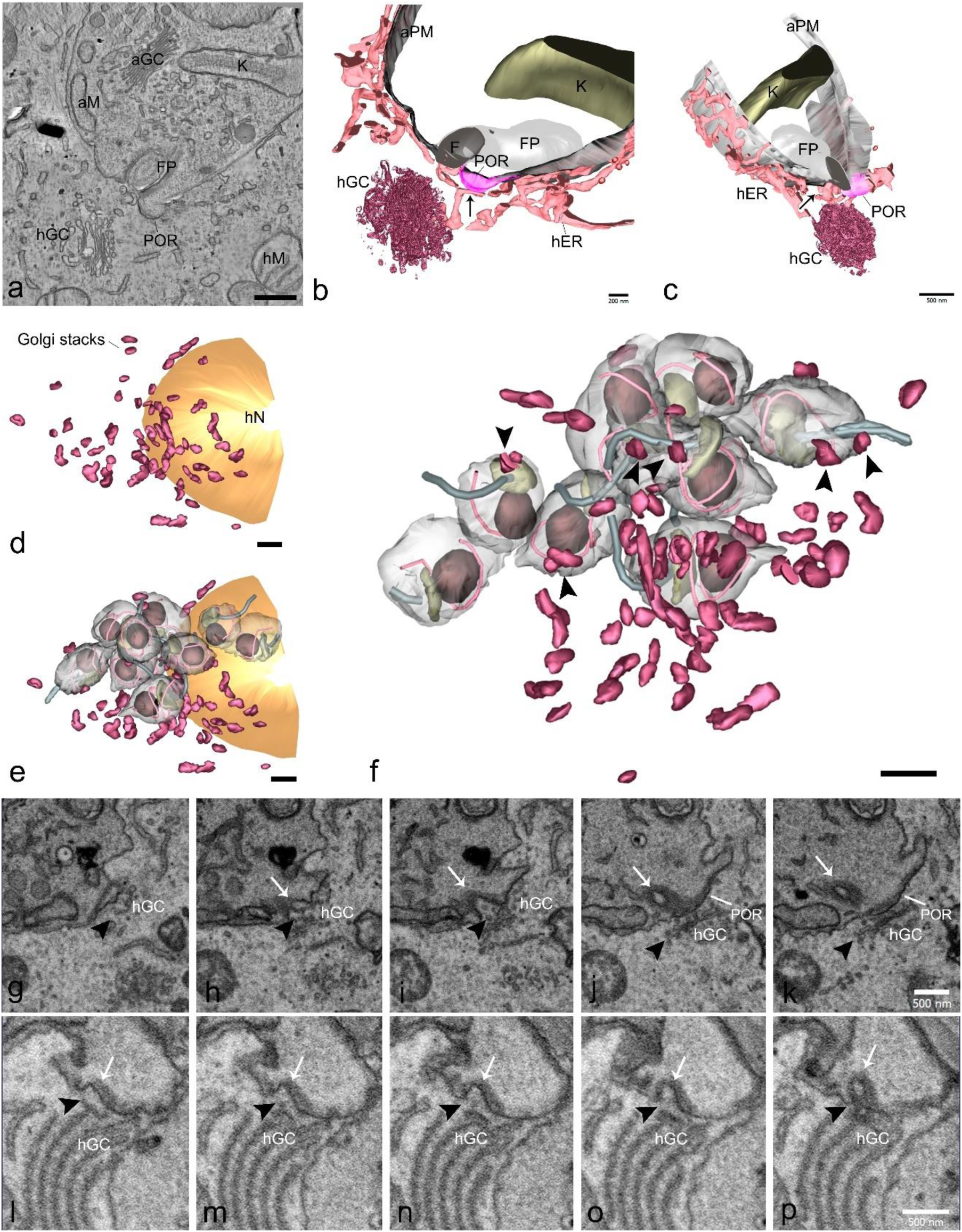
Host Golgi complex fragments during infection. HFF1 cells were infected with trypomastigotes and incubated with the growth medium supplemented with 10% of complete FBS (control) for 48h. Cells were then fixed and processed for TEM. (a) Virtual slice of serial dual-axis tomogram showing the proximity of a Golgi stack (hGC) to the anterior region of the amastigote, close to the preoral ridge (POR) and flagellar pocket (FP). (b,c) Different views of the 3D model produced from the serial tomogram are shown in a. (b) Host ER (hER, pink) surrounds the amastigote membrane (aPM, gray) along the entire volume. Host Golgi was placed close to the amastigote POR and FP. An ER profile placed close to the Golgi could be seen touching the POR membrane domain (arrow). (c) ER, tethered the amastigote membrane along all reconstructed volumes. An ER profile could be seen close to the FP (arrow). (d-p) A slice-and-view series was obtained in a FIB-SEM. (d-f) 3D reconstruction of the volume analyzed showing Golgi stacks (magenta) dispersed through the host cell cytoplasm (d). Golgi stacks were found scattered among the amastigotes (e,f). Golgi stacks get close to the amastigote membrane, especially at the anterior region of the parasite (arrowheads in f). (g-p) Sequential images of the FIB-SEM series showing the region of contact of 2 different amastigotes (g-k and l-p) with a host Golgi stack (hGC, arrowheads). The cytostome is pointed by the arrow. Note the presence of Golgi-derived vesicles at the entrance of the cytostome (arrowheads in h, i). Bars: a,c,g-p – 500 nm; d-f - 2 µm.

Despite the high resolution offered by electron tomography, the cellular volume that can be assessed by serial electron tomography is still limited. Therefore, we used FIB-SEM to acquire an automated serial section from preparations of infected cells 48hpi incubated in a culture medium supplemented with 10% FBS. The reconstructed volume was 6000 µm3. In Figs. 7d and 7e the tridimensional reconstruction shows an overview of the Golgi stacks distribution in infected cells, corroborating our findings in the fluorescence microscopy concerning the fragmentation of the Golgi ribbon and dispersion of the Golgi stacks. The Golgi stacks were distributed among the amastigotes. Moreover, we observed that the proximity of Golgi elements with the amastigote membrane was found only at the anterior region of the parasite, close to the cytostome and the flagellar pocket (Fig 7f). Despite the low resolution of the FIB-SEM technique, the volume imaged allowed us to analyze the ultrastructure of the interaction between the amastigotes and the host organelles and snapshot images that suggests internalization of host vesicles derived from Golgi or ER through the cytostome-cytopharynx complex (Figs 7g-p).

## Discussion

Previous work had investigated the distribution of host cell organelles at the initial steps of *T. cruzi* infection and parasitophorous vacuole formation in non-phagocytic [24] and phagocytic cells [25], and evidenced the essential role of lysosomes, Golgi complex and mitochondria. After escaping to the cytosol and differentiating, amastigotes were registered at the host perinuclear region in several host cell types [26]. In that situation of initial infection amastigotes were found surrounded by endoplasmic reticulum and mitochondria. The authors chose to investigate the relationship with the mitochondria, finding contact sites with the parasite flagellar tip that they hypothesized to function as sensors [26].

Understanding the mechanisms by which *T. cruzi* amastigotes scavenge nutrients from the host cell is important to unveil the factors that can lead to disease progression, particularly in the chronic phase, and for the development of strategies that can inhibit parasite replication.

Sterols are essential lipids in the cellular membranes. They can have several important functions that include the maintenance of the liquid-solid phase of the membranes, allowing for signaling and membrane traffic inside the cells, as well as energy source and reservoir. Different types of sterols can be synthesized among living organisms. *Trypanosoma cruzi*, like other trypanosomatids and fungi, mainly produces ergostane-type sterols [27], which differs from its vertebrate host, in which cholesterol is the major sterol. Because of that, the biosynthetic pathway for the production of ergosterol has been explored as a chemotherapy target for drug development [28].

Although *T. cruzi* synthesize its own sterols, it has been shown that epimastigotes are able to capture cholesterol from the extracellular medium, through endocytosis of LDL [29]. The endocytosed cholesterol can be distributed throughout the cell membranes of the parasite and also stored as cholesterol esters in lipid droplets or lipid inclusions inside reservosomes [12–14]. The main carbon source used by *T. cruzi* for energy production is still a matter of debate. However, some data from the literature points to the important role of lipids as energy fuel [17].

Amastigotes grow and replicate inside the cytosol of host vertebrate cells. Differently from other intracellular protists, such as Leishmania sp. and *Toxoplasma gondii*, that proliferate inside a parasitophorous vacuole [30], amastigotes are free in the cytosol, having direct access to the host cytosol macromolecules and organelles. In this context, it has been shown that amastigotes possess more than 80% in weight of its sterol as cholesterol [15], suggesting that it may scavenge this lipid from the host cell.

In this work, we demonstrated, for the first time, the ability of amastigotes to capture cholesterol from the host cell. We used a fluorescent cholesterol analog, TopFluor Cholesterol (or BODIPY-cholesterol), whose biophysical properties were shown to be very similar to the cholesterol, especially for lipid traffic assays [31]. For that, we tested different methods of delivering TopFChol to cells, to study the impact of the traffic dynamics to the parasite.

The observation that cholesterol was internalized by amastigotes faster when TopFChol was added directly to the growth medium when compared with TopFChol provided inside LDL particles, suggested that amastigotes intercept cholesterol along the pathway from the host cell plasma membrane to intracellular membranes, prior to its passage through the host’s endocytic route. The different dynamics for cholesterol internalization were described in models of lysosomal cholesterol accumulation, using NPC1-deficient fibroblasts [31]. They showed that, when incubated in the medium, TopFChol took more than 18h to start to accumulate in lysosomes while when provided inside LDL particles, it took just 2h. This suggests that in the case of amastigotes, access to TopFChol may happen downstream of lysosomal export. In this context, the role of the host ER emerges as an important factor in this traffic. This is not a surprise since ER is the site of cholesterol synthesis and a major regulator of cholesterol homeostasis in the cell. Previous work estimated that approximately 30% of the LDL-derived cholesterol is exported to the ER without traffic through the plasma membrane [32].

Cholesterol signals in amastigotes were characterized by punctual labeling along the anterior/posterior region of the parasite, with major concentration at the anterior region, close to the flagellar pocket. In most trypanosomatids of medical importance, the flagellar pocket is the sole site of all endo-exocytosis [33]. However, *T. cruzi* possesses an additional structure related to endocytosis, the cytostome-cytopharynx complex [7]. This long membrane invagination opens at the cell surface, close to the flagellar pocket, and is separated from it by a highly glycosylated membrane domain called Preoral Ridge. We have shown that the preoral ridge domain extends from inside the flagellar pocket toward the cytostome and that it may participate in the transport of endocytic cargo that binds to the flagellar pocket membrane to the cytostome where it can be endocytosed [7]. The concentration of cholesterol signal at the anterior region of the amastigotes could indicate the participation of the endocytic portals in its internalization. Indeed, immunogold labeling of infected cells incubated with TopFChol showed strong labeling for the cholesterol at the preoral ridge region, which corroborates our hypothesis. Moreover, the winding pattern of cholesterol labeling inside amastigotes was similar to that already shown for endocytosed transferrin incubated with amastigotes released from host cells, with punctate labels arranged in a helical shape [6], that corresponded to the endocytic tracer passing through the cytopharynx.

Cholesterol intracellular transport in mammalian and yeast cells, where it is best characterized, possesses two main mechanisms: vesicular traffic and non-vesicular traffic. Non-vesicular lipid transport is mediated by membrane contact sites (MCS), regions of close membrane apposition (5-30 nm) between neighboring organelles [34], and accounts for a fast route of lipid exchange. The lipid traffic between membranes is mediated by lipid transfer proteins (LTPs) that can act either as shuttles or bridges, helping to tether membranes and facilitates lipid transfer [35]. EM has been considered the “gold standard” method for the visualization and characterization of MCSs [36]. For that, we resorted to the use of electron tomography (ET) and FIB-SEM to characterize the mechanism of cholesterol uptake by amastigotes. Our results indicate the participation of MCS between ER and amastigotes as a probable mechanism of cholesterol traffic between the host cell and the parasite. Multiples bridges between the ER and the amastigote membrane were seen by ET whose dimensions are compatible with what was previously described for these bridges in intraorganellar MCS.

Besides ER, the host Golgi complex seems also to contribute to the traffic of cholesterol to amastigotes. We observed Golgi ribbon fragmentation and stack dispersion upon infection with *T. cruzi,* highlighting a remodeling of the organelle never explored before in *T. cruzi*-host interaction. Golgi fragmentation is also observed in other pathogenic diseases, the induction of Golgi fragmentation seems to be actively mediated by the pathogens [37]. In *Toxoplasma gondii* infection, Golgi fragmentation and dispersal around the parasitophorous vacuole is important for the acquisition of host sphingolipids by the parasite, by the sequestering of host Golgi-derived vesicles [38]. By analyzing a big volume from an infected cell, we could see the approximation of the host Golgi stacks with the anterior region of amastigotes. The observation of Golgi-derived vesicles close to the entrance of the cytostome suggests that endocytosis of Golgi vesicles could occur. This could explain some previous data from the literature that showed uptake of host TGF-β by amastigotes infecting cardiomyocytes [39]. In this work, Waghabi and colleagues observed, by immunostaining host TGF-β, labeling inside amastigotes but not in trypomastigotes. TGF-β is a cytokine that is secreted by the cells. This way, its productions follow the classical biosynthetic-secretory pathway of the cell, being translated in ER, transferred to Golgi, and secreted to the plasma membrane via Golgi-secreted vesicles. In that work, the major question that remained was how amastigotes had access to this cytokine if it is contained inside secretory vesicles and one of the hypotheses was the ability of amastigotes to endocytose host secretory vesicles. Corroborating for that, we also showed the presence of ER-luminal proteins and Golgi proteins inside amastigotes not only by antibody staining of these proteins but also by using expression vectors to produce tagged proteins.

Although this work does not identify the proteins responsible for the cholesterol traffic into ER-amastigotes MCS, we brought important mechanistic insights into the pathways that *T. cruzi* amastigotes can use to scavenge host molecules. From here, we can now have a clue about the proteins that may participate in these MCS, especially proteins derived from the parasite. It is important to point out that, *T. cruzi* may have lipid transfer proteins localized at its plasma membrane which can deal with cholesterol scavenging from host ER and Golgi. Of note, from what we know about *T. cruzi* cellular physiology, the cholesterol traffic from the parasite cell surface to the cytosol must be dependent on the parasite endocytic pathway, since subpellicular microtubules preclude the contact of any intracellular organelle with the parasite plasma membrane [40]. This way, cholesterol inserted into the membrane should be delivered to the cytostome-cytopharynx complex by lateral diffusion and internalized by endocytosis, after vesicle budding from the cytopharynx.

The relevance of cholesterol scavenging from host became clear with our demonstration that host cholesterol availability directly impaired amastigote growth and development. The presence of cholesterol in the medium was essential to support amastigote growth. Incubation with delipidated FBS decreased amastigote development suggesting that *de novo* synthesis of cholesterol by the host cell was not sufficient to fulfill the sterol requirements for amastigotes. Addition of TopFChol directly to the culture medium was able to restore amastigote grown to levels like the control (cells supplemented with 10% FBS). The total cholesterol concentration in FBS ranges from 300-500 µg/ml [41]. That means the concentration of total cholesterol in the culture medium supplemented with 10% FBS was around 30-50 µg/ml. In our assay we used 100 µg/ml of TopFChol to supplement the delipidated FBS, which could explain not only the restauration of the amastigote grown but also the slightly higher grow when compared with the control.

The relevance of cholesterol scavenging by the amastigotes is still a matter of debate in the literature. Chemotherapy studies focusing on the effect of inhibitors of the sterol biosynthesis pathway may have paved some clues to this question. It is now established in the literature that inhibitors of lanosterol-14a-demethylase (CYP51), are unable to deliver sterile cure in the clinic [42,43] and in animal models [44], probably by the decrease in amastigote grown in tissues and then a lesser necessity for sterol synthesis de novo. In this context, cholesterol may serve as a substitute for endogenous sterols. In epimastigotes, knockout of CYP51 and squalene epoxidase (SQLE) leads to ablation of all endogenous sterols and the only sterol detected was exogenous cholesterol [45]. Although parasites grew slower when compared to wild type, they were viable, suggesting that exogenous cholesterol was sufficient to fulfill the parasite’s sterol requirement, not only metabolically but structurally also.

Fig 8 summarizes our results and hypotheses. We suggest that extracellular-derived cholesterol may have different traffic pathways to amastigotes: [1] cholesterol loaded in LDL-particles are captured by receptor-mediated endocytosis [1a], passing through the early endosomes and late endosomes [1b] being released in the lysosomes [1c], from where it can be transferred to ER and then to amastigotes via MCSs [1d] or it can be transferred directly from lysosomes to amastigotes via MCSs [1e], although this last seems not to be a prominent pathway. [2] Cholesterol in excess in the host cell PM is sequestered in the ER by ER-PM MCSs [2a], which then can be accessed by amastigotes via ER-amastigotes MCSs [2b]. Cholesterol in the ER can be transferred to fragmented Golgi stacks via vesicular transport or from MCSs, and Golgi-amastigotes MCSs may be responsible for cholesterol transfer to amastigotes [2c]. Another alternative is the direct endocytosis of Golgi-derived vesicles by amastigotes [2d]. In this regard, Golgi fragmentation during infection and proximity from amastigoteś endocytic sites may aid this process.

**Fig 8.**
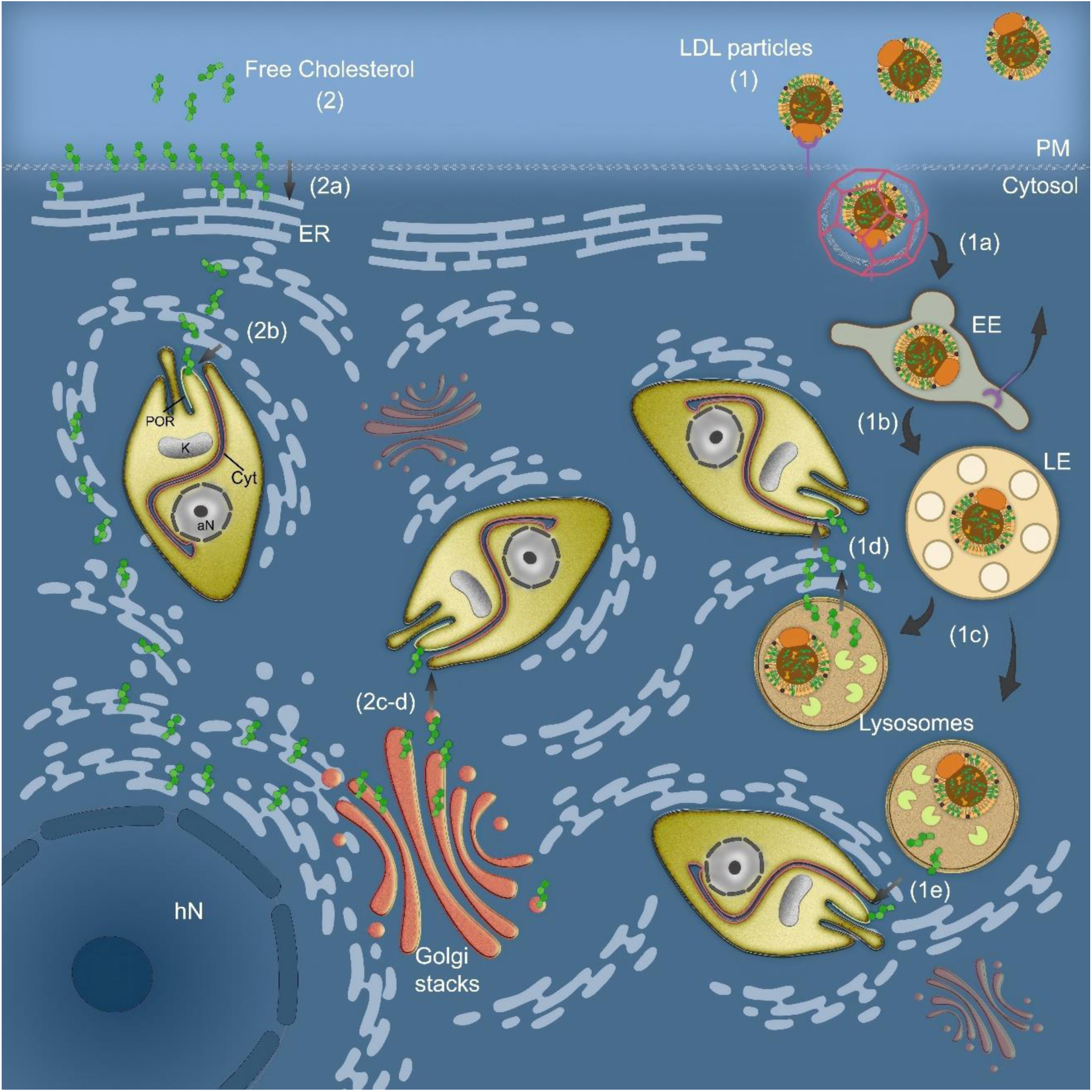
Schematic representation of the routes of cholesterol transport to amastigotes. (1) cholesterol loaded in LDL particles is captured by receptor-mediated endocytosis (1a), passing through the early endosomes and late endosomes (1b) and being released in the lysosomes (1c), from where it can be transferred to ER and then to amastigotes via MCSs (1d) or it can be transferred directly from lysosomes to amastigotes via MCSs (1e). (2) Cholesterol in excess in the PM is equilibrate by ER-PM MCS establishment (2a), which then can be accessed by amastigotes via ER-amastigotes MCSs (2b). Cholesterol in excess in the ER can be transferred to fragmented Golgi stacks via vesicular transport or from MCSs, and Golgi-amastigotes MCSs may be responsible for cholesterol transfer to amastigotes (2c). Another alternative is the direct endocytosis of Golgi-derived vesicles by amastigotes (2d). Amastigote cytostome-cytopharynx complex (Cyt) may concentrate shuttled cholesterol and internalize it through endocytosis. POR (preoral ridge), K (kinetoplast), aN (amastigote nucleus), and hN (host nucleus).

## Materials and Methods

### Cell Culture and Parasites

Human Foreskin Fibroblasts (HFF-1, SCRC-1041, ATCC) were maintained as subconfluent monolayers in Dulbecco’s modified Eagle’s medium (DMEM, Gibco, Grand Island, NY) supplemented with 10% fetal bovine serum (FBS, Gibco, Grand Island, NY), 2 mM L-alanyl-L-glutamine dipeptide in 0.85% NaCl (Glutamax, Gibco, Grand Island, NY) and 1% penicillin-streptomycin (Gibco), at 37°C under 5% CO^2^. Tissue culture Trypanosoma cruzi trypomastigotes from clone Dm28c, were used to infect HFF1 cells with a MOI of 10:1. The development of intracellular amastigotes was followed over time.

### Serum delipidation

FBS was delipidated according to Cham and Knowles (1976) [46]. The delipidated fetal bovine serum (dFBS) was sterilized by filtration with a 0.22 µm membrane (Millex-GV, Millipore S.A., Molsheim, France).

### TopFluor Cholesterol preparation and complexation with LDL particles

TopFluor Cholesterol (Avanti Polar Lipids, catalog number 810255) was dissolved in pure ethanol at a concentration of 1mg/mL. Low-density lipoprotein (LDL) was purified from fresh human plasma as described by Chapman et al. (1981) [47] with some modifications. 100 µg of TopFChol was mixed with 12 mL of human cell-free plasma followed by KBr addition to adjust the density to 1.3 g/mL. The plasma was added to a centrifuge tube with 20 mL of saline solution (150 mM NaCl + 1 mM EDTA). This material was ultracentrifuged at 150.000 *g* in a vertical angle Beckman VTi 50 rotor (Beckman Coulter Inc, Fullerton, CA, USA) at 4 °C for 12 h. The LDL fraction was localized and removed. KBr was added again to the LDL fractions to adjust the density to 1.2 g/cm3, and the material was ultracentrifuged at 150.000 *g* in the same rotor at 4 °C for more 12h. The purified lipoproteins complexed to TopFChol were extensively dialyzed against PBS with 1 mM EDTA.

### TopFChol assays in infected cells

For experiments where TopFChol was added directly to the culture medium, cells were infected with trypomastigotes (MOI 10:1) in a medium containing 10% FBS for 4 hours. Cells were washed to remove parasites that did not enter, and incubation with TopFChol was made 24h later. TopFChol, at a final concentration of 10µM, was diluted in a culture medium containing 10% dFBS, before incubation with cells. For incubations with TopFChol loaded into LDL particles, infected cells were prepared as described above. LDL-TopFChol, at a concentration of 100 µg/ml, was added to the culture medium containing 10% dFBS. At the end of the incubation period, cells were washed with PBS and fixed with 4% (v/v) methanol-free formaldehyde in PBS.

### Quantification of TopFChol internalization by amastigotes

Incubation of infected cells with LDL-TopFChol or free TopFChol were performed as described above. Samples were imaged in in a Zeiss Elyra PS.1 confocal laser microscope equipped with an ACS APO 63.0×1.40 OIL DIC objective. Lasers lines 405, 488, 543, and 633 were used. Z-stack series were acquired with the Zen software (Zeiss) with a frame averaging of 2 and a step size of 0.3 μm. For quantification of amastigote internalization of TopFChol, amastigotes were divided into two halves (Fig. 3a), and labeling was classified based on the presence of the tracer at amastigote’s cell anterior, anterior and posterior or no labeling. Two hundred amastigotes were counted for each condition (from two independent experiments). Statistical analysis was performed by two-way analysis of variance (ANOVA) followed by Bonferronís multiple comparison test, using the Graph Pad Prism 9.5.1 software (La Jolla, CA, USA). . All quantification data represent mean and standard deviation values. Results were considered statistically significant when p < 0.05.

### Amastigote proliferation

HFF cells were infected with trypomastigote (MOI 10:1), incubated for 4h, washed to remove parasites that do not entered cells and incubated for more 20h in medium supplemented with 10% FBS. After, the medium was changed and cells were incubated in medium containing 10% FBS (control), 10% dFBS, or 10% dFBS + 10µM of TopFChol. Cells were fixed 24h later (48hpi) with 4% (v/v) formaldehyde in PBS, pH 7.2 and stained with DAPI. Three hundred host cells were counted for each condition (two independent experiments in triplicate) using Cell Counter plugin of ImageJ. Parasite/cell and percentage of infection (% infection) were normalized by the control sample. Graph plotting and result analysis performed using the One-way ANOVA with Post Hoc Holm-Šídák’s multiple comparisons test, as adequate. The analyses were carried out using the GraphPad Prism 9.5.1 software (La Jolla, CA, USA).

### Fluorescence Microscopy

Infected or non-infected HFF1 cultures, incubated or not with TopFChol, were fixed with 4% (v/v) formaldehyde in PBS, pH 7.2. For immunolabeling experiments, cells were washed in PBS pH 7.2 to remove the fixative, incubated with 150 mM ammonium chloride for 15 minutes, and incubated with a blocking solution (3% BSA, 0.1% saponin in PBS pH 8) for 1h. Primary antibodies were diluted in the blocking solution and incubated for 1h, followed by incubation with secondary antibodies, also diluted in the blocking solution, and incubated for 1h. Samples were DAPI (4′, 6-diamidino-2-phenylindole; Thermo Fisher) stained and mounted on microscopy slides using Prolong Diamond antifade Mountant (Thermo Fisher). Primary antibodies used were: anti-PDI (Thermo Fisher – PA582640); anti-TGN-38 (Sigma Aldrich – T9826); anti-Lamp1 (Thermo Fisher – PA1654A); and anti-Ssp4 (monoclonal 2C2; [48]), kindly provided by Dr. Renato Mortara (Unifesp, São Paulo, Brazil). Secondary antibodies used were Alexa Fluor 594 Goat Anti-Rabbit or Anti-Mouse IgG (Thermo Fisher), and Alexa Fluor 647 Goat Anti-Mouse IgG (Thermo Fisher). Alternatively, cells were incubated with Cell Light ER-RFP or Cell Light Golgi-RFP reagents (Thermo Fisher) overnight, according to the manufacturer’s instructions, washed in sterile PBS, and then incubated with TopFChol, with the same conditions described above, if that was the case. Cells were fixed as described above.

All samples were imaged in a Zeiss Elyra PS.1 confocal laser microscope equipped with an ACS APO 63.0×1.40 OIL DIC objective. Lasers lines 405, 488, 543, and 633 were used. Z-stack series were acquired with the Zen software (Zeiss) with a frame averaging of 2 and a step size of 0.3 μm.

### Sample preparation for transmission electron microscopy (TEM)

Infected or non-infected cultures were fixed by using 2.5% (v/v) glutaraldehyde in 0.1 M cacodylate buffer, pH 7.2, for 1 h at room temperature, post-fixed using an osmium-thiocarbohydrazide-osmium (OTO) protocol [7,49], dehydrated in acetone series and embedded in epoxy resin. Samples were cut in ultrathin sections in a Leica EM UC7 Ultramicrotome and stained post-embedding with 5% (w/v) uranyl acetate and lead citrate. Samples were imaged in a Tecnai Spirit electron microscope (Thermo Fisher) operating at 120 kV, or in Hitachi HT 7800 (Hitachi) operating at 120 kV.

### Immunogold labeling

Infected or non-infected cultures incubated with TopFChol for 24h as described above, were fixed with 4% (v/v) methanol-free formaldehyde, 0.2% (v/v) glutaraldehyde in 0.1 M cacodylate buffer, pH 7.2, for 1 h at room temperature. Samples were dehydrated in ethanol series, infiltrated, and embedded in LR White acrylic resin. Samples were cut in ultrathin sections in a Leica EM UC7 Ultramicrotome and sections were collected in nickel grids. Grids were blocked with a blocking buffer (1.5% BSA in PBS pH 8) for 1h, then incubated with BODIPY FL Polyclonal Antibody (Thermo Fisher) diluted 1:100 in the blocking buffer for 1h, and with secondary antibody goat anti-rabbit IgG Nanogold (Thermo Fisher) diluted 1:100 in blocking buffer for 1h. After several washes in water, nanogold labeling was enhanced using Pierce Silver Stain Kit (Thermo Fisher), according to the manufacturer’s instructions. Grids were not post-stained and were observed in a Hitachi HT 7800 operating at 120 kV.

### Electron tomography

Samples processed for TEM were cut in 200-nm-thick serial sections using a Leica EM UC7 Ultramicrotome. Sections were collected onto formvar coated copper slot grids and post-embedded stained as described before. Single or Dual-axis tilt series (±65° with 1° increment) were produced in a Tecnai Spirit electron microscope (Thermo Fisher) operating at 120 kV and coupled to a 2k x 2k pixel CCD camera. Tilt series were reconstructed by weight back projection using ETOMO software from the IMOD package [50].

### Focused Ion Beam-Scanning Electron Microscopy

Samples prepared for TEM were trimmed and glued to an SEM stub using carbon tape and metalized with gold. Samples were imaged using an Auriga dual-beam microscope (Zeiss) equipped with a gallium-ion source for focused-ion-beam milling, a field-emission gun, and an in-lens secondary electron detector, for SEM imaging. The cross-sectional cut was made at ion beam currents of 2.0 uA and an accelerating voltage of 30 kV. Backscattered electron images were recorded at an accelerating voltage of 1.8 kV and a beam current of 0.8 nA, in the immersion lens mode, using a CBS (Concentric Back Scatter) detector. A series of backscattered electron images were recorded in “slice-and-view” mode, at a magnification of 15 K, with a pixel size of 3.5 nm and milling step size of 30 nm. After image capture, back-scattered electron images had their contrast inverted, to resemble conventional TEM images. FIB-SEM serial images were aligned using ETOMO software from the IMOD package.

### Tridimensional Reconstruction

Serial tomograms and serial sections obtained in FIB-SEM were segmented and rendered using 3DMOD from the IMOD package.

## Acknowledgments

The authors thank to Mayara Menezes and Msc. Veronica dos Santos, for the maintenance of the cell cultures and infection protocols, to Heloá Estevam for the help with serum delipidation and LDL isolation. A special acknowledgment to Msc. Azuil Barrinha for the valuable insights about data statistics analysis. We thank to the Centro Nacional de Biologia Estrutural e Bioimagem (CENABIO) – Advanced Microscopy Unit, for the use of the Transmission Electron Microscopes and FIB-SEM.

## Supporting Information

**S1 Fig.**
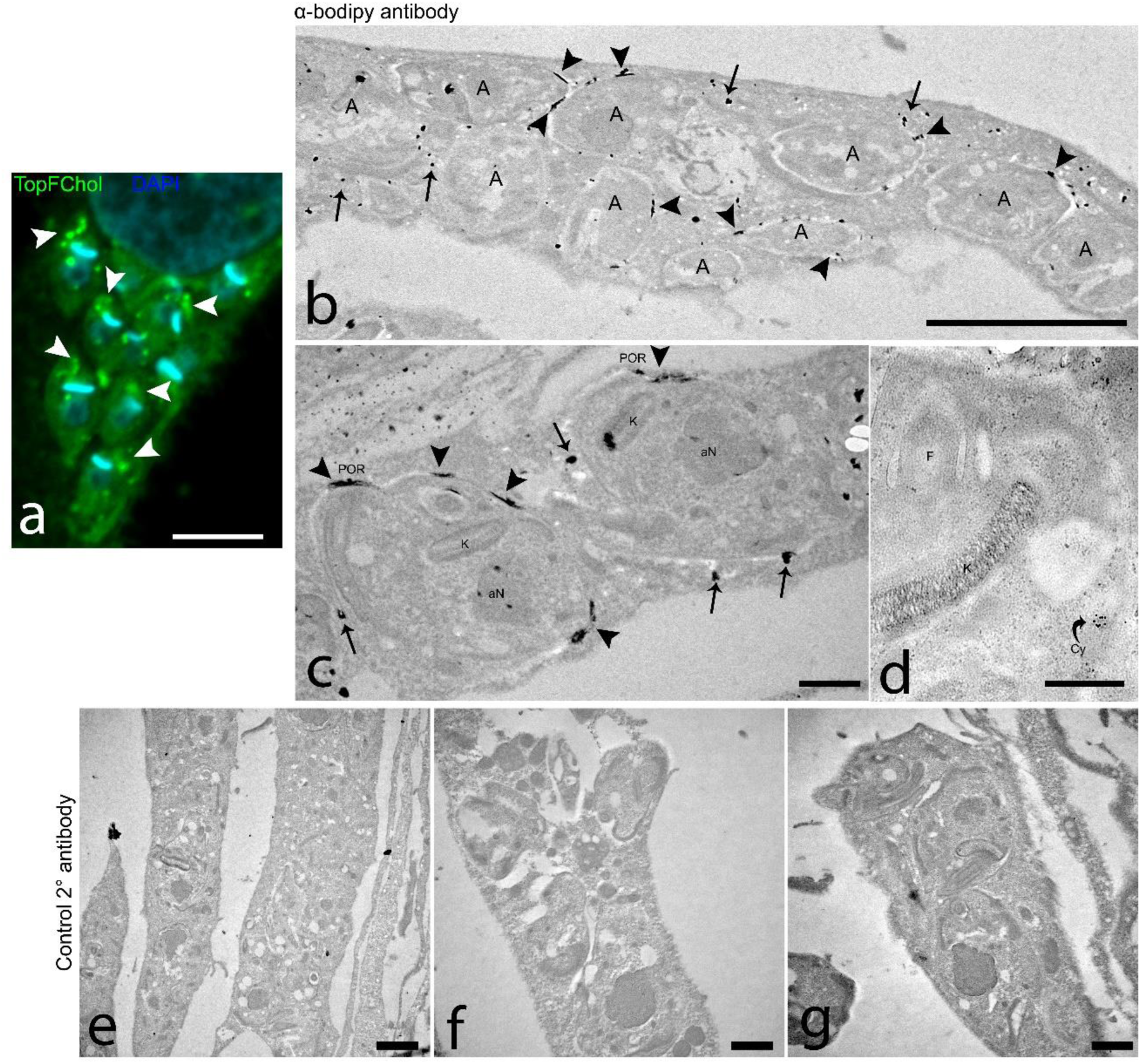
The pattern of TopFChol inside amastigotes. HFF1 cells were infected with trypomastigotes and, 24h after, TopFChol was added directly to the growth medium supplemented with 10% of dFBS. (a) Single-plane fluorescence image from the z-stack confocal series of infected cells incubated 24h with TopFChol. Note that the TopFChol signal (green) showed a strong concentration at the amastigote anterior region, close to the kinetoplast (bar shape in blue) (arrowheads). The labeling followed a curved path towards the cell posterior. (b-d) Infected cells, incubated with TopFChol for 24h were processed for TEM using acrylic resin and immunogold labeled with a primary antibody anti-bodipy moiety of TopFChol and reaction was silver enhanced. Grids were not post-stained. (b) Low-magnification image showing an overview of an infected cell. An intense labeling reaction was observed at puncta around the amastigotes plasma membrane (arrowheads) as well as in regions of the host cytosol (arrows). (C) The strong electron-dense labeling was observed at the anterior region of the amastigote, specifically at the membrane domain of the preoral ridge (POR) (arrowheads). (d) Some gold particles can be seen at the membrane of the cytopharynx (Cy). (e-g) Images of the same material incubated only with the secondary antibody and silver stained shows the absence of reaction. Bars: a,b – 5 µm; c,f,g - 1 µm; d-500 nm; e – 2 µm.

**S2 Fig.**
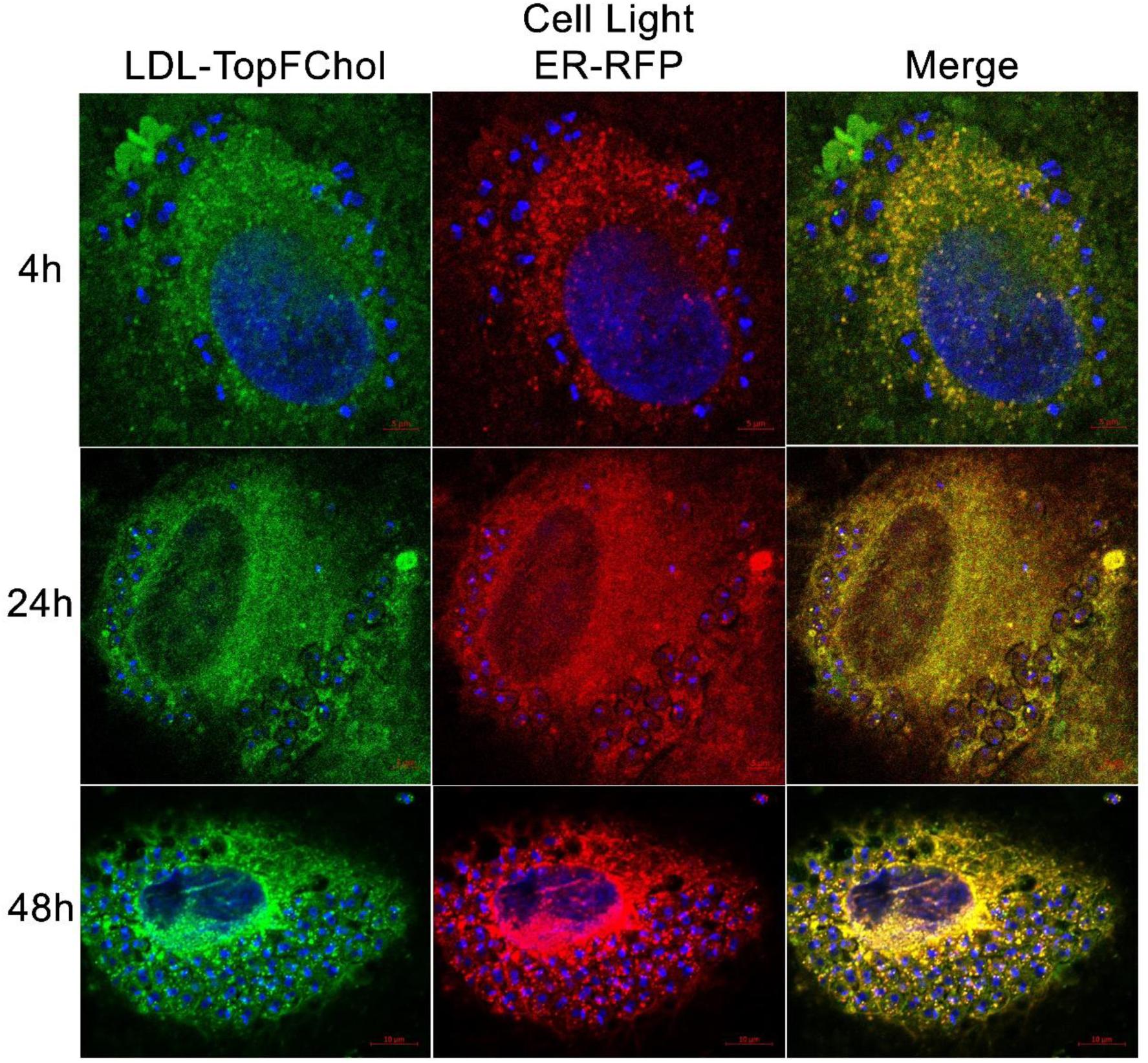
Dynamics of TopFChol internalization by intracellular amastigotes through incubation with LDL-TopFChol and Cell Light ER-RFP. HFF1 cells were infected with trypomastigotes and, 24h after, incubated with Cell Light ER-RFP for 24h. LDL-TopFChol was then added to a growth medium supplemented with 10% dFBS and incubated for 4h, 24h, and 48h. Single-plane fluorescence images from the z-stack confocal series are shown. Colocalization of the TopFchol signal and Cell Light ER-RFP was observed inside amastigotes at all time points. At 48h, many colocalizing punctual labeling was observed. Bars: 5 µm

**S3 Fig.**
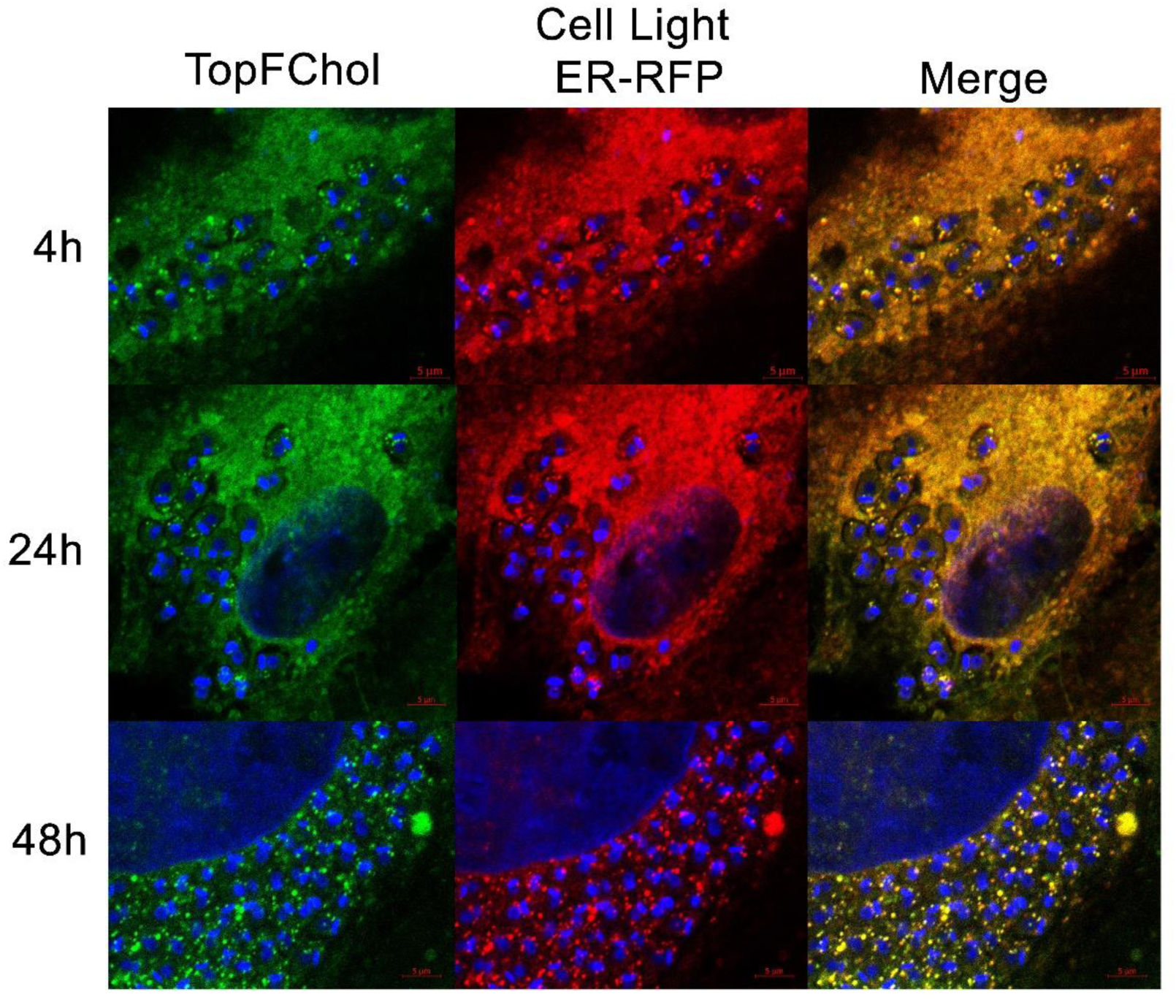
Dynamics of TopFChol internalization by intracellular amastigotes through incubation with TopFChol directly to the culture medium and Cell Light ER-RFP. HFF1 cells were infected with trypomastigotes and, 24h after, incubated with Cell Light ER-RFP for 24h. TopFChol was then added to a growth medium supplemented with 10% of dFBS and incubated for 4h, 24h, and 48h. Single-plane fluorescence images from the z-stack confocal series are shown. TopFChol labeling inside amastigotes has been registered since 4h of incubation. TopFChol signal colocalized with Cell Light ER-RFP inside amastigotes at all time points. At 48h, many colocalizing punctual labeling was observed. Bars: 5 µm

**S4 Fig.**
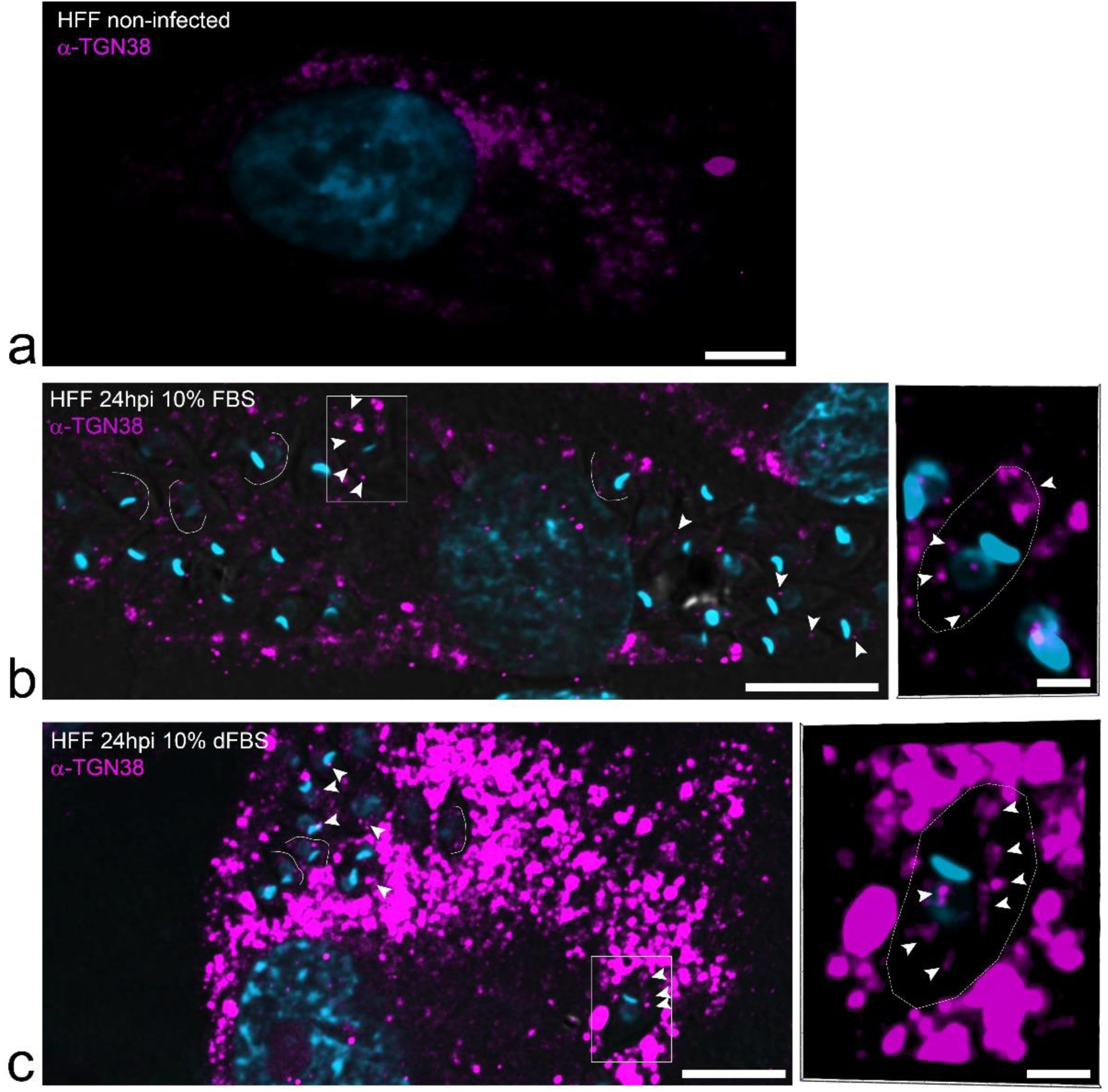
Golgi dynamics is altered during infection. Non-Infected or infected cells incubated in different conditions of lipid availability were fixed and immunolabelled with antibody anti-TGN-38, to mark host Golgi complex. (a) Non-infected cell shows continuous labeling (magenta) restricted to the perinuclear region. (b) Infected cell incubated with complete serum 24hpi shows dispersed punctual labeling of Golgi complex. Labeling for TGN-38 could be seen inside amastigotes along an arched path from the amastigote anterior to posterior (white lines). At the rectangle area, seen in high magnification at the inset, TGN-38 labeling marks the FP and POR region in amastigotes and several other punctual labelings from anterior to the post-nuclear region of the parasite (arrowheads). (c) In infected cells incubated with delipidated serum, Golgi labeling was also punctate and dispersed through the host cell cytoplasm, however, labeling was more intense. It was also possible to observe TGN-38 labeling inside amastigotes along an arched path from the amastigote anterior to posterior (white lines). Labeling was also seen in punctual structures inside the amastigotes (inset from the rectangle region, arrowheads). DAPI (blue) stained DNA-containing structures. Bars: 10 µm; insets – 2 µm.

**S1 Movie. Virtual sections of a dual-axis serial tomogram of HFF1 cells 48hpi.** Cells were grown in a culture medium supplemented with 10% of complete FBS (control). The tomogram shows an area of close contact between host ER and Golgi stack with the anterior region of amastigote membrane. Available in https://figshare.com/account/items/26195582/edit

